# ASIP promoter variants predict sesame coat color in Shiba Inu dogs

**DOI:** 10.1101/2021.10.01.462523

**Authors:** S.N Belyakin, D.A. Maksimov, M.A. Pobedintseva, P.P. Laktionov, D. Voronova

## Abstract

Different patterns of coat color pigmentation in dogs are produced by a sophisticated interaction of several genes. Understanding the mechanisms underlying the diversity of coat colors and their inheritance is important for professional breeders because it helps to predict the phenotypes of the progeny. Although genetics of the main coat colors in dogs is extensively studied, there are types of coat pigmentation that are not explained yet. Recently a new model connected the variants in ASIP gene promoters with different coat colors in dogs. Here we used this model as a framework to investigate the genetics of the rare sesame coat color in Shiba Inu dogs. We determined the combination of two alleles of ASIP gene that determine sesame coat color. This finding can be used by the breeders to produce the dogs with this rare coat color pattern. We also demonstrate the incomplete dominance between the ASIP alleles involved in sesame coat formation. These results are in good agreement with the new model explaining how different levels of ASIP gene expression affect the regulation of pigment synthesis in melanocytes.

## Introduction

Genetics of coat colors in domestic dogs is an example of extensively characterized biological system with peculiar genetic interactions including several epistases (Kaelin and Barsh, 2013; Schmutz and Berryere, 2007). Diversity of coat colors in dogs results from different combinations of two pigments, dark eumelanin and yellow/red pheomelanin, that are synthesized in melanocytes and loaded into the growing hair (Ando et al., 2012; Prota, 1980; Tobin, 2008).

Agouti Signaling Protein produced by *ASIP* gene switches the type of pigment that is produced in melanocytes and thus plays central role in coat color formation (Barsh, 2006). ASIP protein is a ligand of Melanocortin 1 Receptor (MC1R) on the surface of melanocyte. In the absence of ASIP protein MC1R promotes eumelanin synthesis. Binding of ASIP to MC1R switches melanocyte to pheomelanin production (Lu et al., 1994; Millar et al., 1995; Ollmann et al., 1998).

In many animal species the concentration of ASIP oscillates in the skin on the dorsal side of the body resulting in banded pattern of pigmentation along the hair (Kaelin and Barsh, 2013; Vrieling et al., 1994). For example, at the starting phase of hair growth the concentration of ASIP in the skin is low and predominantly eumelanin is produced in melanocytes and loaded to the hair tip. At some point ASIP concentration increases and melanocytes are switched to pheomelanin synthesis resulting in red pigmentation as the hair continues to grow. When the amount of ASIP goes down again a new dark band may appear in the hair.

In domestic dogs alleles of *ASIP* gene are known to produce a number of different patterns. When other genes participating in coat color formation are intact, dominant allele *Ay* leads to a solid red coat pattern. There are variants that produce black-and-tan or saddle tan patterns. Most recessive allele (allele *a*) forms the solid black coat color (Willis, 1989). The wild type allele *aw* forms the wolf-sable coat color characterized by a banded pattern of hair pigmentation due to ASIP oscillation.

Different alleles of *ASIP* gene were previously associated with particular DNA polymorphisms. *Ay* allele correlated with the double mutation c.(244G>T;248G>A) in *ASIP* gene (Berryere et al., 2005). Allele *a* was determined as a missense mutation *ASIP*: c.286C>T (Kerns et al., 2004). Allele *at* was assigned to a SINE insertion in the intron 1 of *ASIP* gene that was found in saddle tan and black-and-tan dogs (Dreger and Schmutz, 2011). These two patterns were further differentiated by a presence of a short duplication in the intron of *RALY* gene (Dreger et al., 2013). This paradigm explains most of the *ASIP* gene effects on coat pigmentation in dogs and corresponding tests are routinely used by breeders. However there are important exceptions that could not be explained by these tests.

Recent study determined that apart from allele *a*, which is indeed the missense loss-of-function mutation, other alleles in fact affect the promoters of *ASIP* gene thus modifying its activity (Bannasch et al., 2021). Two *ASIP* gene promoters are known in many animals (Vrieling et al., 1994). The Ventral Promoter (VP) predominantly activates *ASIP* gene on the ventral part of the body. Its activity gradually decreases towards the dorsal surface. Transcription from VP is permanent, so the coat color on the ventral surface of the body is normally red due to pheomelanin deposition. Another promoter, which was called Hair Cycle Promoter (HCP) regulates *ASIP* gene activity mostly on the dorsal surface of the body. The activity of this promoter normally circulates leading to a banded hair pattern (Kaelin and Barsh, 2013; Vrieling et al., 1994).

As it was recently shown there are at least two different variants of VP and five variants of HCP. The combinations of these variants accurately determine different alleles of *ASIP* gene (Bannasch et al., 2021). The strong promoter VP1 paired with strong promoter HCP1 corresponds to *Ay* allele. In this case the concentration of ASIP protein is permanently high in all parts of the skin which leads to the uniform red pigmentation of hairs. The double mutation *ASIP*: c.(244G>T;248G>A) that was previously considered as a causative for *Ay* allele is in fact a linked marker of the promoter combination VP1+HCP1(Bannasch et al., 2021).

A weaker variant VP2, which is actually wild type, together with strong HCP1 variant produces predominantly red pigmentation but on the dorsal surface of the body hairs have dark tips due to eumelanin produced at the beginning of hair growth. Here we call this combination allele *Ays*, a shaded version of *Ay*. This allele was previously genetically undistinguishable from *Ay* because it is also linked to the double mutation *ASIP*: c.(244G>T;248G>A) (Bannasch et al., 2021).

Two wild type variants, VP2 and HCP2 produce wild type (wolf sable) pattern characterized by banded hair pigmentation typical to wolves (Bannasch et al., 2021). This combination that corresponds to allele *aw* and can be now determined directly by PCR test.

Three HCP variants (HCP3-5) have lost their ability to activate *ASIP* gene on the dorsal surface of the body. In combination with VP2 promoter these HCP variants produce black and tan coat pattern characterized by black coat on the dorsal surface and red pigmentation on the ventral surface (Bannasch et al., 2021). This variant corresponds to allele *at*.

Accordingly a combination of stronger VP1 promoter with HCP4 (allele *asa*) is responsible for saddle tan pattern, which differs from black and tan pattern by a wider expansion of red pigmentation, including the head (Bannasch et al., 2021). Interestingly, no other combinations of VP1 with non-functional variants of HCP were reported.

The hierarchy of *ASIP* gene alleles was established as follows: *Ay*>*Ays*>*aw*>*asa*=*at*>*a*. Two alleles, *asa* and *at*, are codominant (Dreger et al., 2013). The dogs with *asa*/*at* genotype show intermediate coat pattern between black and tan and saddle tan. Black pigmentation on their heads forms a characteristic “widow’s peak”-shaped pattern.

In Shiba Inu dogs there is a specific coat pattern called sesame (Figure 1). This pattern resembles the black and tan with the solid black areas replaced by the black overlay. According to FCI breed standard, there are different subtypes of sesame classified as sesame, black sesame or red sesame. These subtypes are differentiated primarily by the amount of black shading.

**Figure 1.**
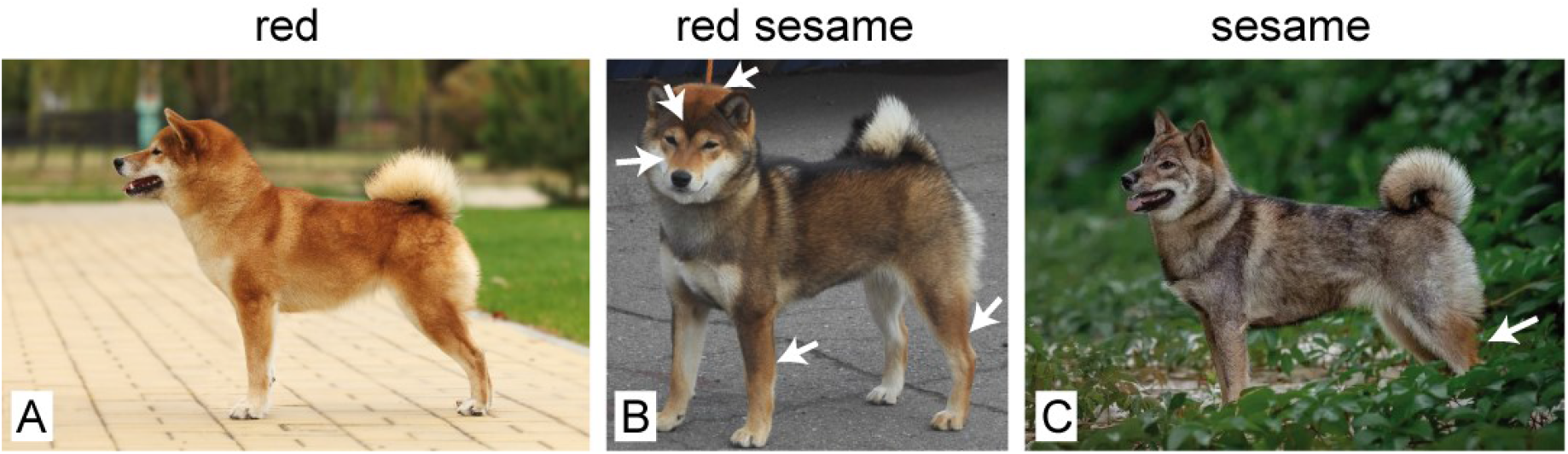
Examples of red (A), red sesame (B) and sesame (C) coat colors in Shiba Inu. Compared to sesame, red sesame dogs have considerably more red pigmentation that is specifically distributed along the body, as indicated with white arrows. A characteristic “widow-peak”-shaped dark pigmentation between the eyes is shown on red sesame dog (B).

Most of the dogs in Shiba Inu breed are red; a few percent are black and tan (also see the statistics in (Dreger et al., 2019)). Sesame coat colors are the rarest ones. The inheritance of sesame coat color is unclear and puzzles the breeders and dog owners for decades. Here we applied the recent findings about *ASIP* gene alleles to investigate the genetics of sesame coat color in Shiba Inu.

## Materials and Methods

### Samples

Shiba Inu buccal swabs were used as a source of genomic DNA for this study. The samples were sent to VetGenomics laboratory by the owners for genetic testing. The dogs used in this study were initially selected using the phenotype information indicated in the Shiba Inu database (http://www.shiba-pedigree.ru/). The owners of all dogs used in this study signed the agreement of participation in the study and sent the photographs of their dogs to confirm the coat color.

### Genotyping of ASIP promoters

Genomic DNAs were isolated from the swabs and used in PCR reactions (using BioMaster HS-Taq PCR-Color (2×) mastermix, Biolabmix, Russian Federation) with the primers for VP and HCP published in Supplementary Table 5 of the previous study (Bannasch et al., 2021). Genotypes were determined by assessing the lengths of PCR products in 1% agarose gel in 1x Tris-Acetate-EDTA buffer.

### Sequencing the MC1R gene

The coding sequence of MC1R gene was amplified using the primers: Forward: 5’-cctcaccaggaacatagcac-3’, Reverse: 5’-ctgagcaagacacctgagag-3’. The 1007 bp PCR product was purified by Polyethylene Glycol (PEG) precipitation (Lis, 1980) to remove primers. The purified PCR product was used for Sanger sequencing on 3500 Genetic Analyser (Applied Biosystems).

## Results

One of the most amazing misfits concerning *ASIP* gene alleles interactions is sesame coat pattern in Shiba Inu dogs. According to the breeders’ reports, sesame dogs have genotypes *aw*/*aw* or *aw*/*at*. Red sesame dogs are *Ay*/*at*, as detected by a widely used genetic test for *ASIP* double mutation c.[244G>T;248G>A] that was previously considered as a causative for *Ay* allele. The problem is that *Ay*/*at* dogs genotyped by this test may be red sesame or red with little if any black shading (Figure 2). The reason for this variation remained elusive.

**Figure 2.**
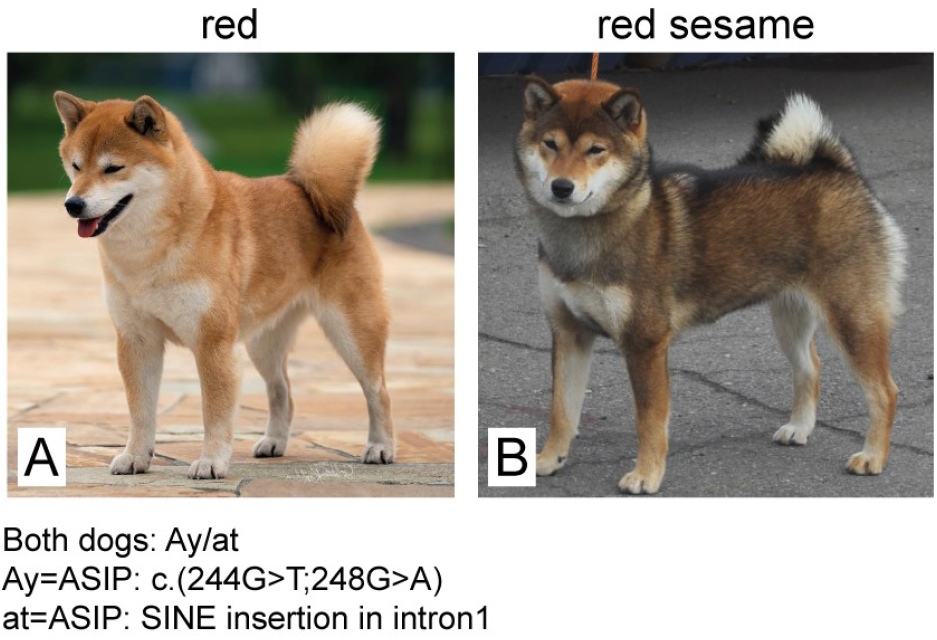
Example of two dogs with *Ay*/*at* genotype, where *ASIP*:c.(244G>T;248G>A) double mutation was as a marker of *Ay* allele. (Bannasch et al., 2021). Both dogs in A and B possess *ASIP*:c.(244G>T;248G>A) mutation but show a striking difference in phenotype. The dog in A is purely red, while the dog in B is red sesame.

To address this question we genotyped promoters of Shiba Inu dogs with red and sesame coat colors. Using PCR approach we determined different VP and HCP variants in 57 dogs living in Russian Federation, including 42 red dogs, and 15 sesame or red sesame dogs. The phenotypes were verified using the photographs from pedigree database (http://www.shiba-pedigree.ru/) as well as pictures provided by the owners.

This analysis provided a clear concordance between coat color and genotype determined using the new findings about *ASIP* gene promoters (Table 1). All the dogs having red coats possessed the most dominant *Ay* allele in their genotypes (*Ay*=VP1+HCP1). Genotyping of the red sesame dogs revealed genotype *Ays*/*at* (*Ays*=VP2+HCP1), which clearly distinguished them from the red *Ay*/*at* dogs. The phenotypically darker sesame dogs were found having the previously reported *aw*/*at* genotype. In these cases *aw* allele was directly determined by VP2+HCP2 promoter combination.

**Table 1.**
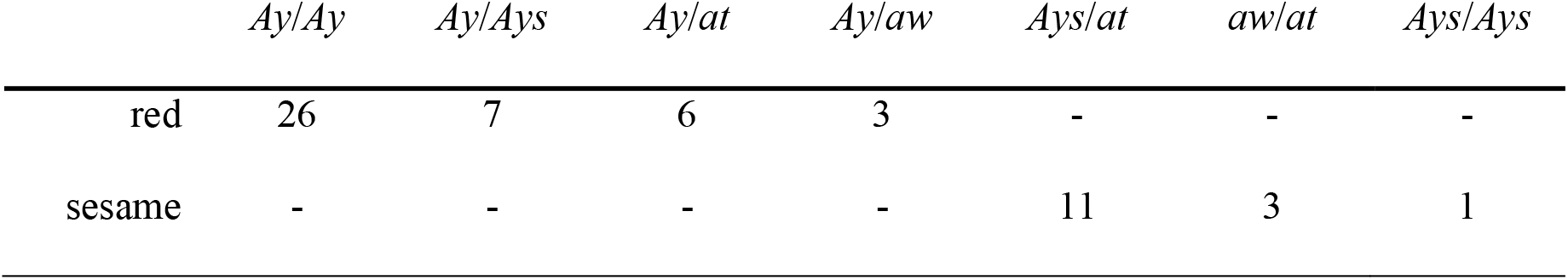
Genotyping *ASIP* gene promoter variants in 57 red and sesame Shiba Inu dogs.

|  | *Ay/Ay* | *Ay/Ays* | *Ay/at* | *Ay/aw* | *Ays/at* | *aw/at* | *Ays/Ays* |
| --- | --- | --- | --- | --- | --- | --- | --- |
| red | 26 | 7 | 6 | 3 | - | - | - |
| sesame | - | - | - | - | 11 | 3 | 1 |

Thus our results demonstrate that red sesame Shiba Inu dogs have genotype *Ays*/*at* in locus A and directly confirm the darker sesame genotype to be *aw*/*at*. No dogs with *aw*/*aw* genotype were found in this study. This observations explains the puzzling inheritance of sesame coat color. To illustrate this we genotyped a family that produced red sesame dog (Figure 3). Both parents were red. Genotyping of *ASIP* gene promoters revealed that Sire has *Ay*/*at* combination while Dam was *Ay*/*Ays*. Their red sesame daughter possesses *Ays*/*at* genotype.

**Figure 3.**
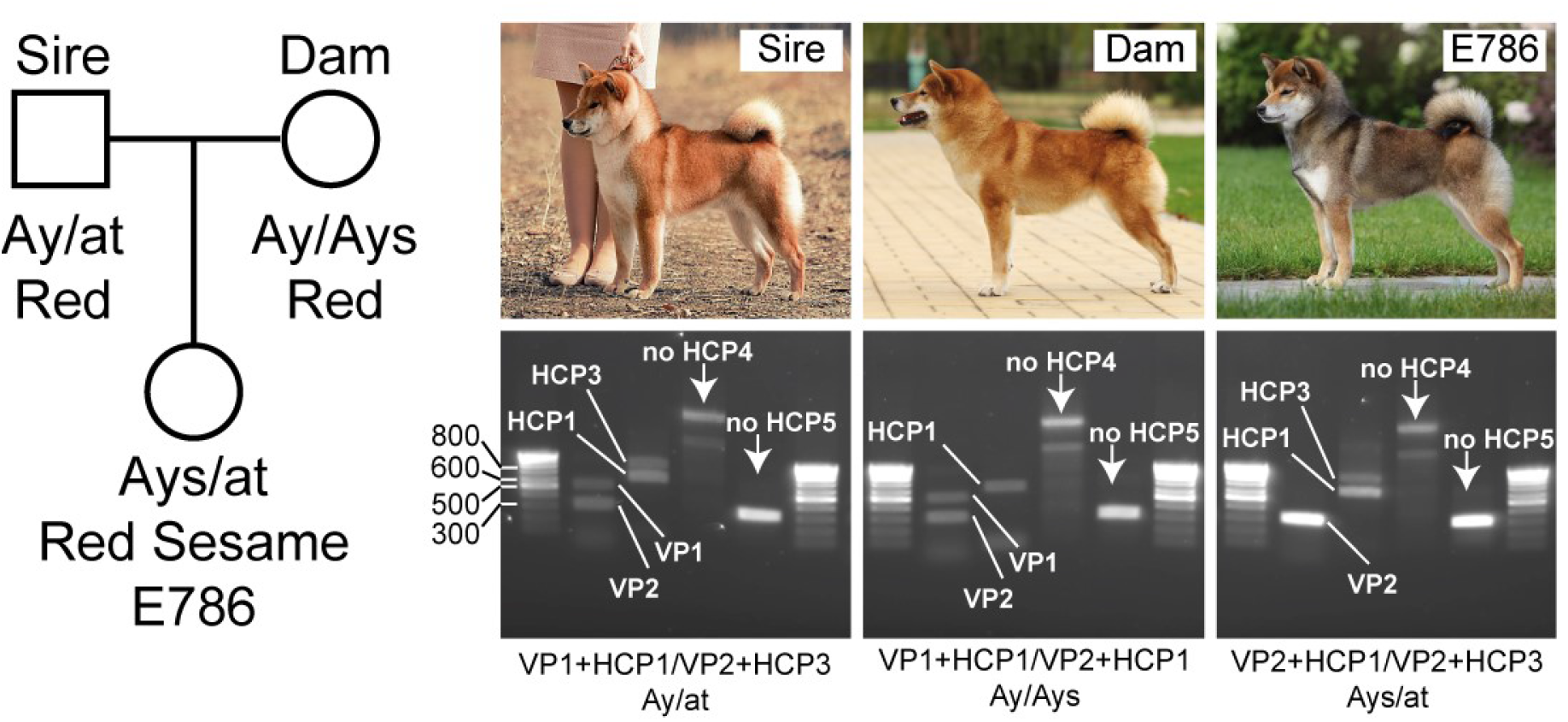
Genotyping of the red sesame dog and her red parents. Red parents and their red sesame daughter were PCR-genotyped for *ASIP* gene promoter variants as described in (Bannasch et al., 2021). The genotyping of the Sire revealed the presence of VP1, VP2, HCP1 and HCP3 promoter variants, which uniquely corresponds to *Ay*/*at* genotype as indicated under the gel photograph (Bannasch et al., 2021). The Dam possesses VP1, VP2 and HCP1 promoters, which correspond to *Ay*/*Ays* genotype. Their daughter inherited *Ays* from Dam and at from Sire as shown by the presence of VP2, HCP1 and HCP3 variants. DNA marker sizes are shown on the firs photograph.

To exclude the possible presence of mutations in MC1R gene that could lead to a shaded coat color we sequenced this gene in these three dogs. No mutations were detected in MC1R gene indicating that locus E does not play role in sesame phenotype formation (data not shown).

Typical red sesame Shiba Inu dogs that were used in this study have *Ays*/*at* genotype. This could indicate that incomplete dominance of the *Ays* allele over the *at* allele exists. Alternatively red sesame coat color would be produced by *Ays* alone and the recessive at allele does not have any impact on hair pigmentation. In the latter case *Ays*/*at* dogs should not be phenotypically different from *Ays*/*Ays*.

*Ays* allele is quite rare in Shiba Inu dogs however we were lucky to find one dog with *Ays*/*Ays* genotype. As shown on the Figure 4, this dog has dark shading that is similar to sesame coat color, but this overlay is generally lighter and much more red pigmentation is seen as compared to *Ays*/*at* dog. This example suggests that *Ays* allele is incompletely dominant over at allele and does not compensate for the lack of HCP activity on the dorsal side of the body.

**Figure 4.**
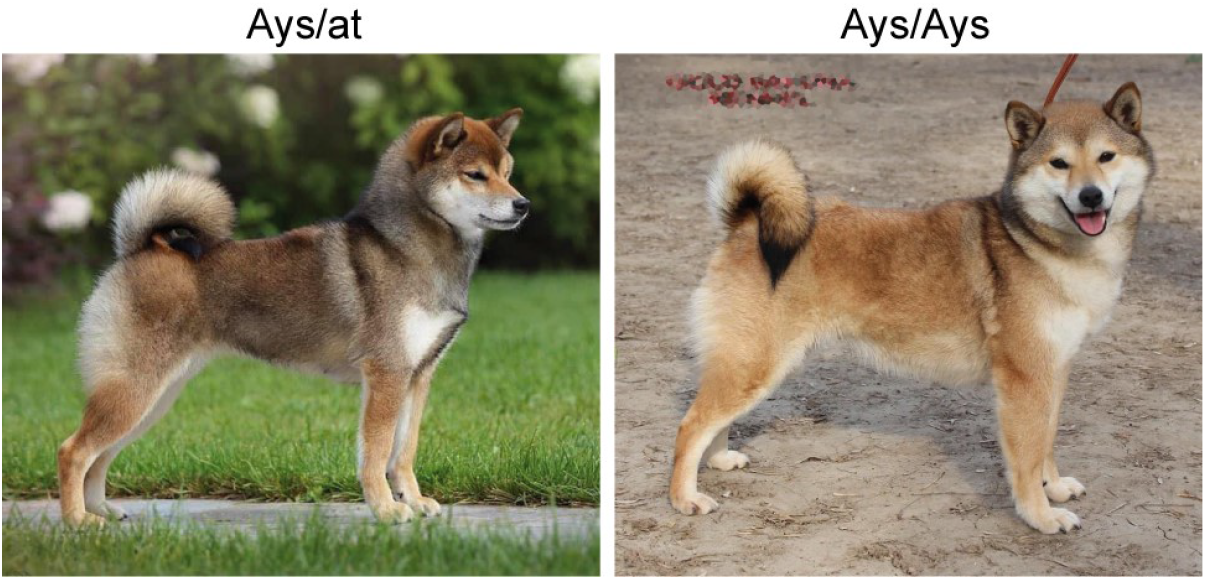
Incomplete dominance of *Ays* allele over *at* allele in red sesame Shiba Inu. *Ays*/*at* dog has a typical red sesame coat. *Ays*/*Ays* dog discovered in this study demonstrates significantly more red in coat color although with obvious dark shading clearly distinguishing it from red Shiba Inu dogs.

## Discussion

Here we tested the recently published model explaining *ASIP* gene effects of the coat pigmentation in dogs (Bannasch et al., 2021). This work allowed us to explain the genetics of red sesame coat color in Shiba Inu dogs, which provides a useful tool to the breeders.

Importantly, our data suggest the incomplete dominance of *Ays* allele over at allele in Shiba Inu. Another example of incomplete dominance between *ASIP* alleles was described in welsh corgi pembroke: *asa*/*at* combination produces “widow-peak”-shaped pattern on the dog head, which is intermediate between saddle tan and black-and-tan (Dreger et al., 2013). Our data suggest that *Ays* allele partially compensates the inability of at allele to produce ASIP protein on the dorsal side of the body in *Ays*/*at* Shiba Inu dogs. It remains unclear whether this effect appears in other dog breeds but to our opinion it is in good concordance with differential promoter activities of *ASIP* alleles (Bannasch et al., 2021).

Our work was focused on the genotyping of *ASIP* gene promoters. This design cannot completely exclude involvement of other genes to sesame coat color. Indeed, for example recently characterized eA allele of MC1R gene showed somewhat similar pattern of pigmentation in different dog breeds (Anderson et al., 2020). As this allele is ancient in canine lineage, it was possible that it exists in Shiba Inu dogs. However direct sequencing of MC1R gene in red sesame dog detected no mutations in its CDS. Together with perfect correlation of red sesame coat color with *Ays*/*at* genotype this suggests that locus E plays no role in sesame coat color.

## Acknowledgements

Authors are grateful to all Shiba Inu breeders and owners who agreed to participate in this study anonymously or publicly, particularly to Elena Gerasimova, Elena Shubnikova, Karine Kaliyan, Evgeniya Makaricheva, Dmitri Mikhailov, Nadezhda Sadovnikova, and Julija Tarhanova.

## Notes

### Competing Interest Statement

The authors have declared no competing interest.

